## Supporting Information for "A unique peptide recognition mechanism by the human relaxin family peptide receptor 4 (RXFP4)"

Brief description of what this file includes:

Supplementary Fig. 1 | Characterization of recombinant INSL5.

Supplementary Fig. 2 | Synthesis and characterization of DC591053.

Supplementary Fig. 3 | Functional validations of the receptor constructs and purification of the complexes.

Supplementary Fig. 4 | Cryo-EM data processing and validation.

Supplementary Fig. 5 | Near-atomic resolution model of the complexes in the cryo-EM density maps.

Supplementary Fig. 6 | Conformational changes upon RXFP4 activation.

Supplementary Fig. 7 | Comparison of the peptide-binding pocket of RXFP4 with other class A GPCRs.

Supplementary Fig. 8 | MD simulations of INSL5-bound active RXFP4.

Supplementary Fig. 9 | MD simulations of INSL5 and its B chain.

Supplementary Fig. 10 | Peptidomimetic agonism and subtype selectivity of compound 4 and DC591053

Supplementary Fig. 11 | Superimposition of INSL5 from the INSL5–RXFP4–$G_i$ complex structure with insulin or IGF-1.

Supplementary Fig. 12 | Superimposition of INSL5 to insulin or IGF-1 in complex with cognate receptors.

Supplementary Table 1 | Cryo-EM data collection, refinement and validation statistics.

Supplementary Table 2 | Interactions of INSL5, compound 4 and DC591053 with RXFP4.

Supplementary Table 3 | Ligands-mediated inhibition of forskolin-induced cAMP accumulation.

Supplementary Table 4 | Effects of residues mutation on ligands-mediated inhibition of forskolin-induced cAMP accumulation.

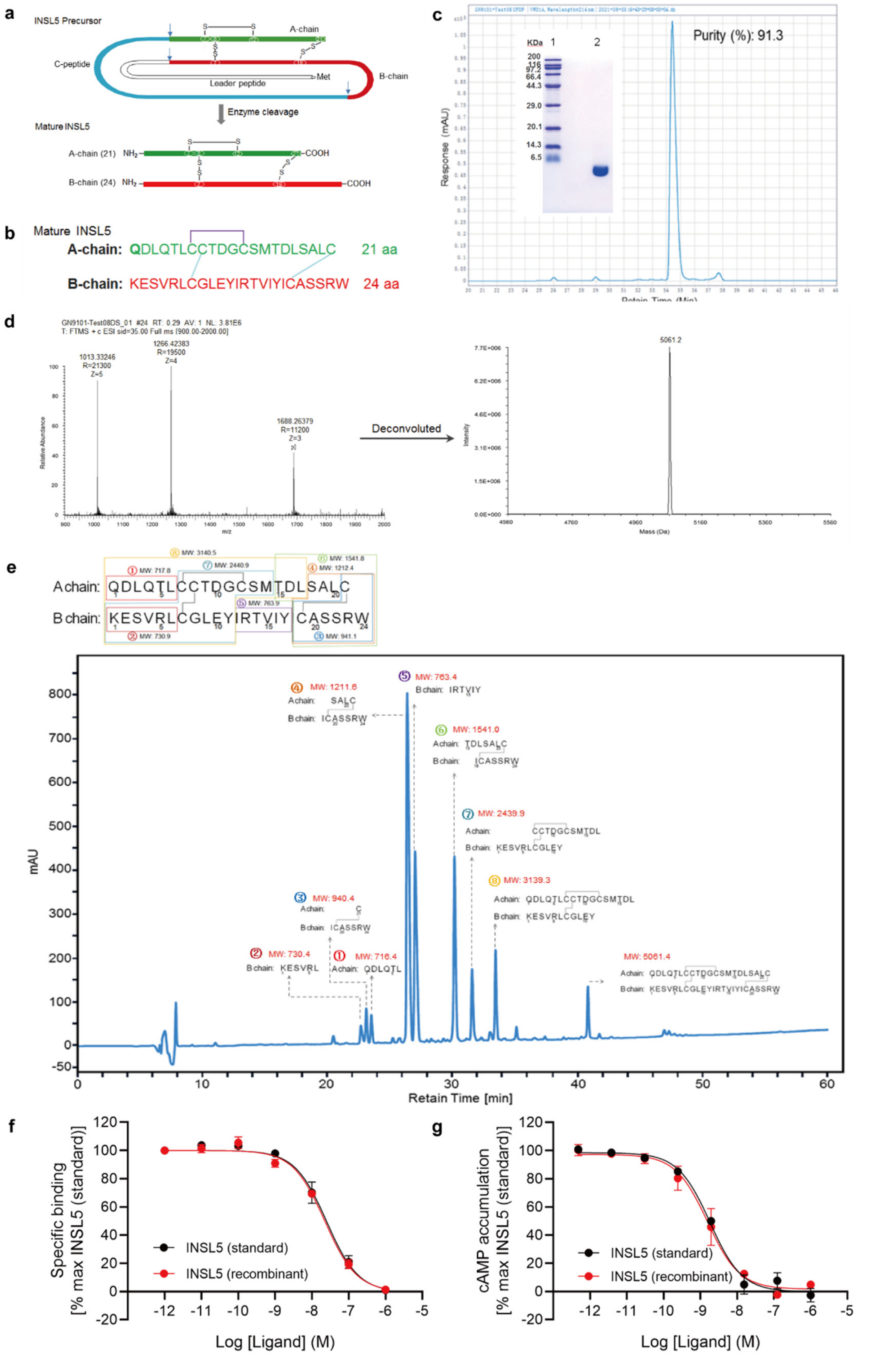

**Supplementary Figure 1. Characterization of recombinant INSL5. a,** The structure of the INSL5 precursor and its conversion to mature INSL5. **b,** Amino acid sequence of the recombinant

INSL5 used in this study. The dash indicates disulfide bond. **c,** Purity analysis of recombinant INSL5 by non-reducing SDS-PAGE and RP-HPLC. Well 1 is the marker, well 2 is a typical batch of recombinant INSL5. The purity of INSL5 in this batch is 91.3%. **d,** Mass spectrometry analysis of INSL5. The measured molecular mass of 5,061.2 Da corresponds to the expected value (5,061.8 Da) of the INSL5 (N-terminal Q of A chain not converted to pE). **e,** Chymotrypsin-generated peptide mapping by LC-LC/MS and assignments for the amino acid sequences of the peptides. Measured molecular mass in red; structure assignment and theoretical molecular weights in black. **f-g,** The recombinant INSL5 was able to bind and activate RXFP4 as determined by europium-labeled ligand competitive binding **(f)** and cAMP accumulation assays **(g)** compared to INSL5 (standard), a control peptide containing native amino acids (N-terminal Q of A chain converted to pE). Source data are provided as a source data file.

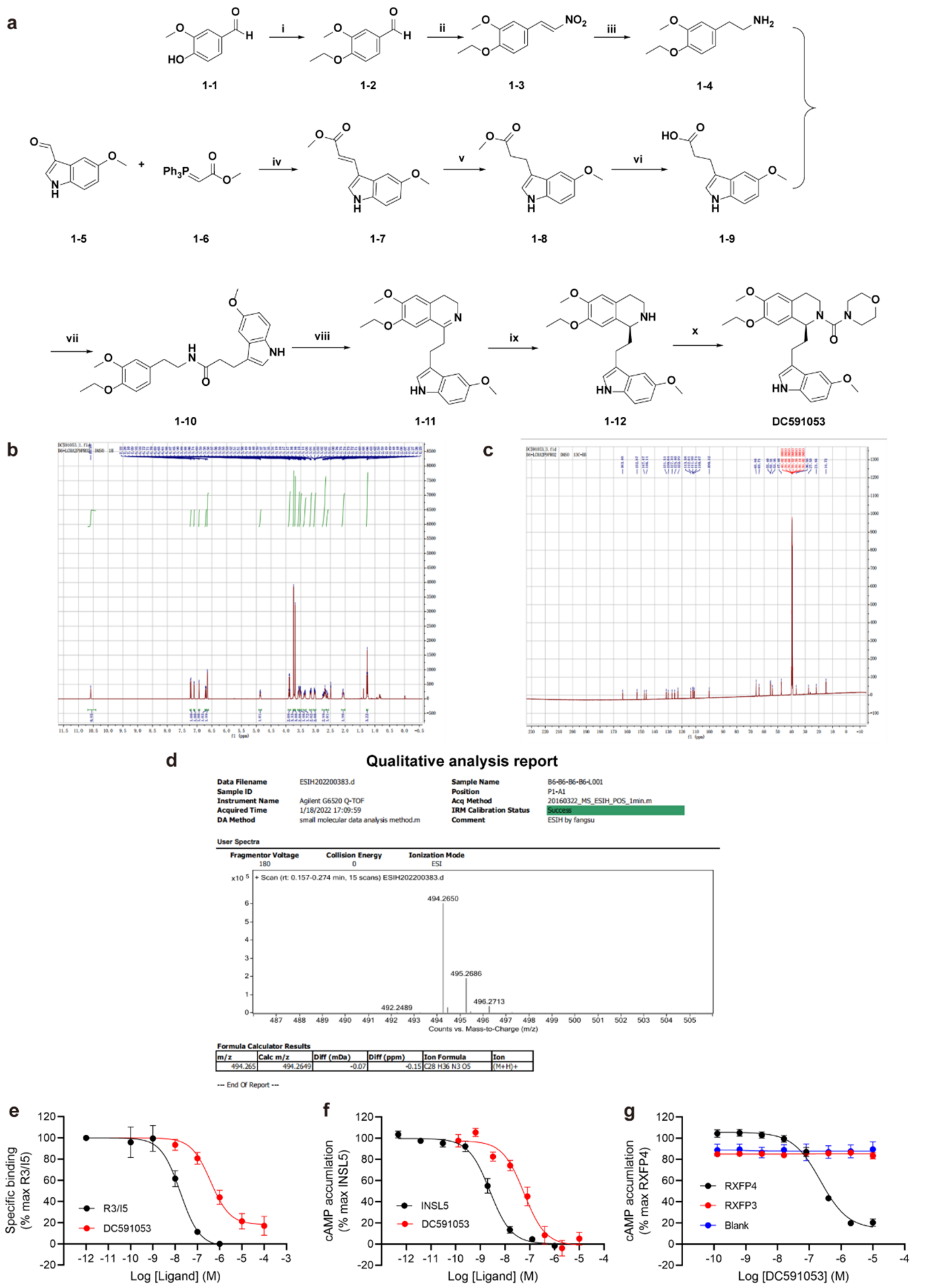

**Supplementary Figure 2. Synthesis and characterization of DC591053. a,** Synthetic route to the compound **DC591053**. (i) $CH_3CH_2I$, $CH_3CN$, $K_2CO_3$, 80 °C, Ar, 90%; (ii) $CH_3COONH_4$, $CH_3NO_2$, 100 °C ,1 h, 65%; (iii) $LiAlH_4$, anhydrous THF, 0 - rt, Ar, 78%; (iv) Toluene, Ar, 110 °C, 16 h, 60%; (v) 10% $Pd(OH)_2/C$, $H_2$, MeOH, 40 °C, 5 h, 95%; (vi) 1M NaOH, MeOH, rt, 93%; (vii) HATU, TEA, rt, DCM, 90%; (viii) $POCl_3$, $CH_3CN$, 80 °C, Ar, 92%; (ix) RuCl[(*R*,*R*)-Tsdpen](*p*-cymene),

$AgSbF_6$, $La(OTf)_3$, HCOONa, $H_2O$/MeOH= 1:1, Ar, rt, 60%; (x) 4-morpholinecarbonyl chloride, DIPEA, DCM, 0 °C to rt, 90%. **b,** $^1$H NMR spectrum of compound DC591053 (500 MHz, DMSO-*d*$_6$). **c,** $^{13}$C NMR spectrum of compound DC591053 (125 MHz, DMSO-*d*$_6$). **d**, HRMS spectrum of compound DC591053**. e-f,** DC591053 was able to bind and activate RXFP4 as determined by europium-labeled ligand competitive binding **(e)** and cAMP accumulation assays **(f)**. **g,** DC591053 neither cross-reacted with RXFP3 nor showed any activity in parental cells. Source data are provided as a source data file.

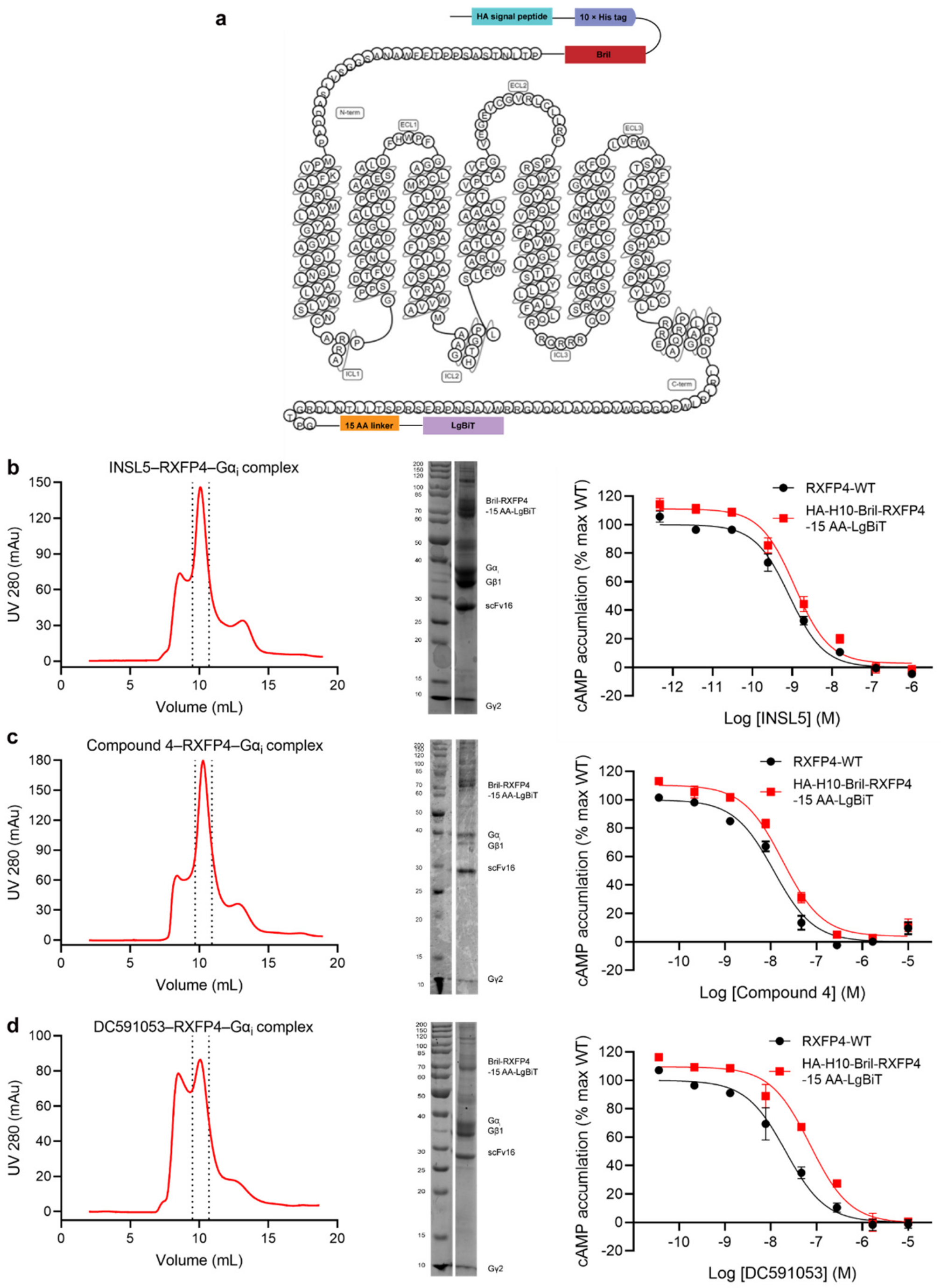

**Supplementary Figure 3. Functional validations of the receptor constructs and purification of the complexes. a**, Schematic diagram of the receptor constructs used for structure determination. **b-d**, Analytical size-exclusion chromatography (left) and SDS-PAGE/Coomassie blue stain (middle) of the purified INSL5–RXFP4–$G_i$ **(b)**, Compound 4–RXFP4–$G_i$ **(c)** and DC591053–RXFP4–$G_i$ **(d)**. The right panel in **b-d** is INSL5, compound 4 and DC591053 induced cAMP accumulation in wild-type (WT) and modified RXFP4 constructs. Data shown were means ± S.E.M. from three independent experiments performed in quadruplicate. Source data are provided as a source data file.

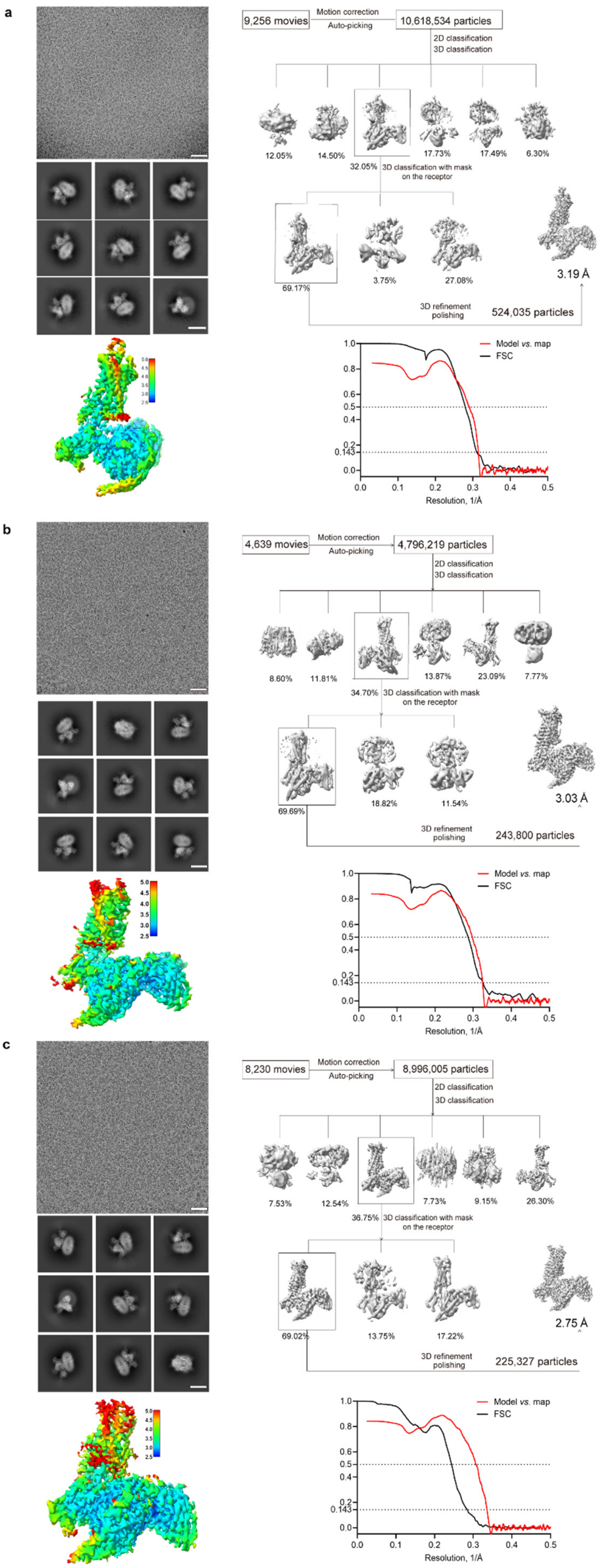
a
9,256 movies
Motion correction
Auto-picking
10,618,534 particles
2D classification
3D classification
12.05%
14.50%
32.05%
17.73%
17.49%
6.30%
3D classification with mask on the receptor
69.17%
3.75%
27.08%
3.19 Å
3D refinement polishing
524,035 particles
Model vs. map
FSC
Resolution, 1/Å
b
4,639 movies
Motion correction
Auto-picking
4,796,219 particles
2D classification
3D classification
8.60%
11.81%
34.70%
13.87%
23.09%
7.77%
3D classification with mask on the receptor
69.69%
18.82%
11.54%
3.03 Å
3D refinement polishing
243,800 particles
Model vs. map
FSC
Resolution, 1/Å
c
8,230 movies
Motion correction
Auto-picking
8,996,005 particles
2D classification
3D classification
7.53%
12.54%
36.75%
7.73%
9.15%
26.30%
3D classification with mask on the receptor
69.02%
13.75%
17.22%
2.75 Å
3D refinement polishing
225,327 particles
Model vs. map
FSC
Resolution, 1/Å

**Supplementary Figure 4. Cryo-EM data processing and validation. a**, INSL5–RXFP4–$G_i$ complex: top left, representative cryo-EM micrograph (scale bar: 40 nm) and two-dimensional (2D) class averages showing distinct secondary structure features from different views (scale bar: 5 nm); top right, flow chart of cryo-EM data processing; bottom left, local resolution distribution map of the complex; bottom right, FSC curves of overall refined receptor. **b,** Compound 4–RXFP4–$G_i$ complex: top left, representative cryo-EM micrograph (scale bar: 40 nm) and 2D class averages showing distinct secondary structure features from different views (scale bar: 5 nm); top right, flow chart of cryo-EM data processing; bottom left, local resolution distribution map of the complex; bottom right, FSC curves of overall refined receptor. **c,** DC591053–RXFP4–$G_i$ complex top left, representative cryo-EM micrograph (scale bar: 40 nm) and 2D class averages showing distinct secondary structure features from different views (scale bar: 5 nm); top right, flow chart of cryo-EM data processing; bottom left, local resolution distribution map of the complex; bottom right, FSC curves of overall refined receptor. These experiments were repeated twice independently with similar results. Source data are provided as a source data file.

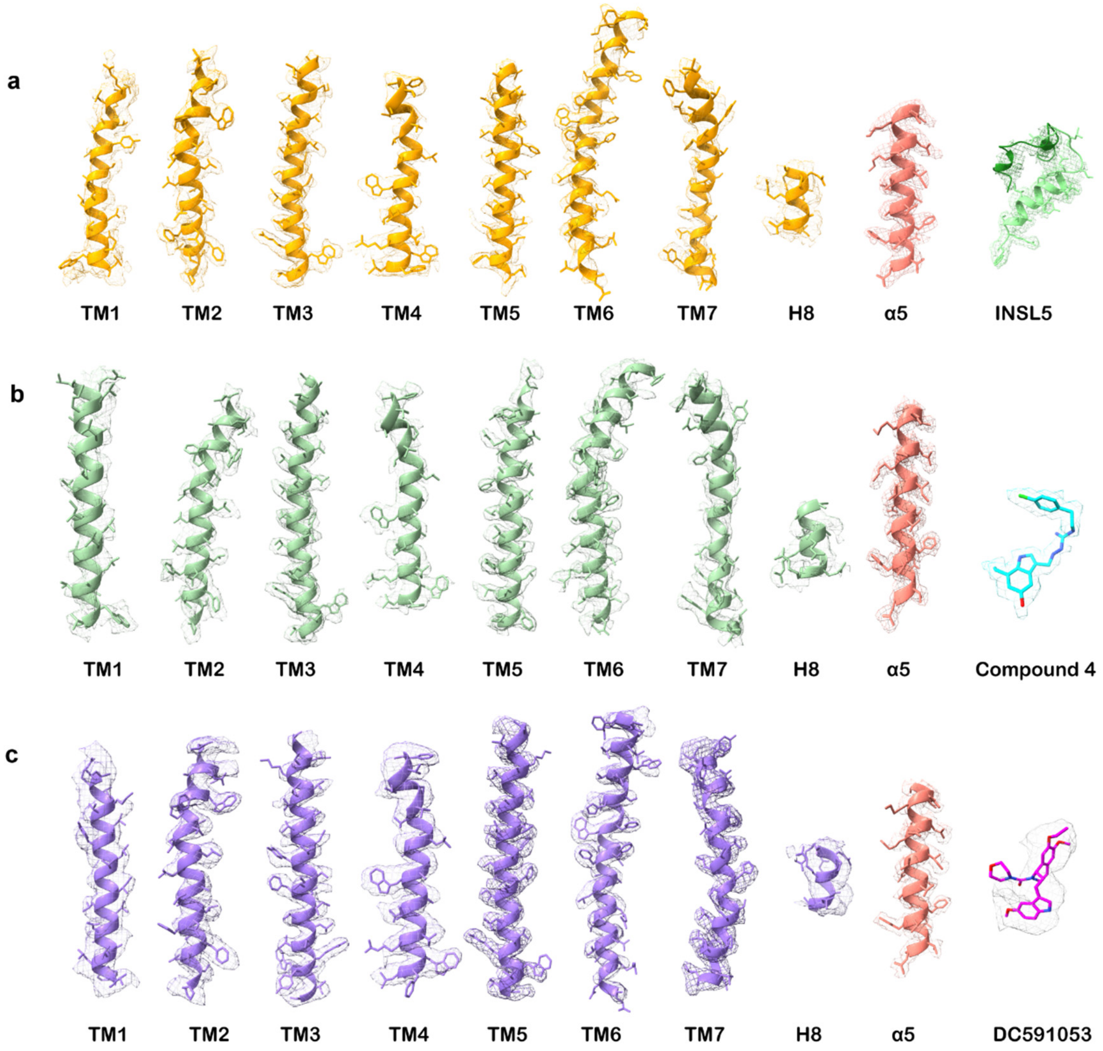

**Supplementary Figure 5. Near-atomic resolution model of the complexes in the cryo-EM density maps. a,** EM density map and model of the INSL5–RXFP4–$G_i$ complex are shown for all seven-transmembrane (7TM) α-helices, helix 8, INSL5 and the α5-helix of the $G_i$ subunit. **b,** EM density map and model of the compound 4–RXFP4–$G_i$ complex are shown for all 7TM α-helices, helix 8 compound 4 and the α5-helix of the $G_i$ subunit. **c,** EM density map and model of the DC591053–RXFP4–$G_i$ complex are shown for all 7TM α-helices, helix 8, DC591053 and the α5-helix of the $G_i$ subunit.

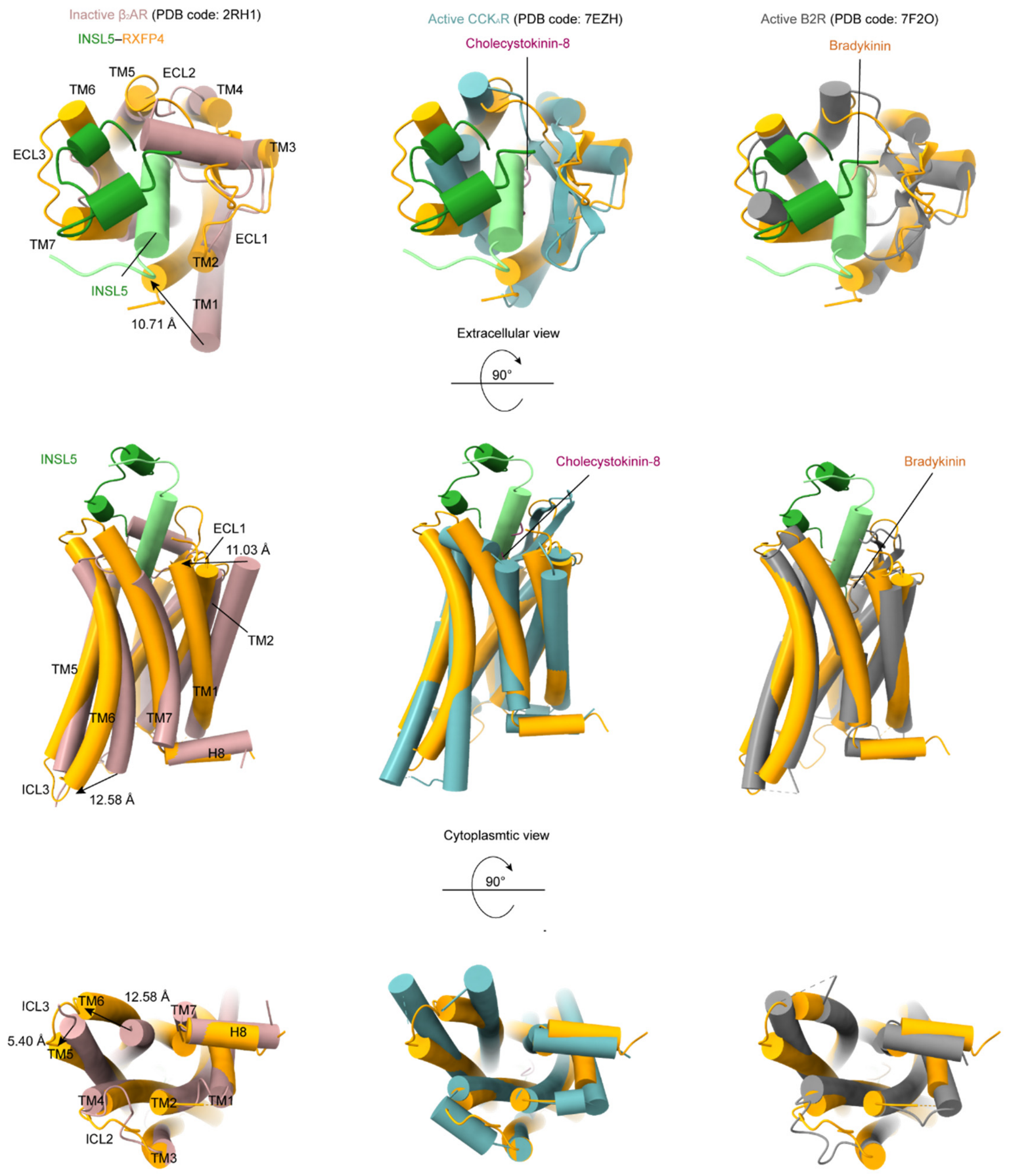

**Supplementary Figure 6. Conformational changes upon RXFP4 activation**. Comparison of active RXFP4 with inactive β2-adrenergic receptor (β2AR) (PDB code: 2RH1)[1] and both agonist-bound and G protein-coupled active cholecystokinin A receptor ($CCK_AR$) (PDB code: 7EZH)[2] and type 2 bradykinin receptor (B2R) (PDB code: 7F2O)[3]. G proteins were omitted for clarity.

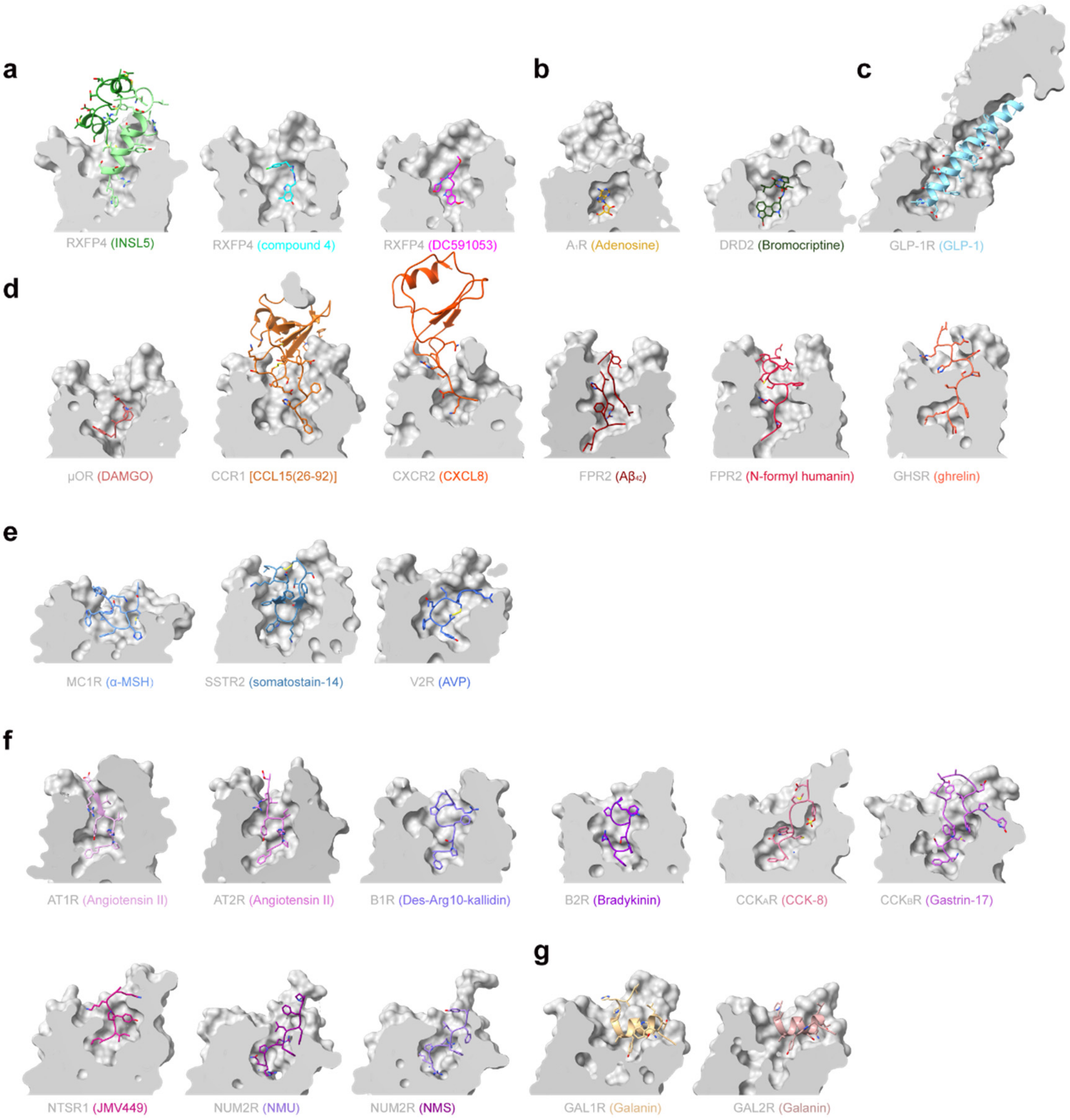

**Supplementary Figure 7. Comparison of the peptide-binding pocket of RXFP4 with other class A GPCRs.** The binding cavity of INSL5, compound 4 and DC591053-bound RXFP4 (**a**) were compared with that of other GPCRs **(b-g)** including small molecule bound receptors [adenosine in adenosine $A_1$ receptor ($A_1R$) (PDB code: 7LD4) and bromocriptine in D2 dopamine receptors (DRD2) (PDB code: 7JVR)] **(b)**, glucagon-like peptide-1 (GLP-1)-bound glucagon-like peptide-1 receptor (GLP-1R) (PDB code: 6X18) **(c)**, the N terminus of peptides inserts into the TMD core such as DAMGO-bound μ-opioid receptor (μOR) (PDB code: 6DDE), C-C chemokine ligand 15 [CCL15(26-92)]-bound C-C chemokine receptor type 1 (CCR1) (PDB code: 7VL9), C-X-C motif chemokine ligand 8 (CXCL8)-bound C-X-C chemokine receptor type 2 (CXCR2) (PDB code: 6LFO), $A\beta_{42}$-bound formyl peptide receptor 2 (FPR2) (PDB code: 7WVY), *N*-formyl humanin-bound FPR2 (PDB code: 7WVX) and ghrelin-bound growth hormone secretagogue receptor (GHSR) (PDB code: 7NA7) **(d)**, the middle segment of cyclic peptides inserts into the TMD binding cavity such as α-melanocyte-stimulating hormone (α-MSH)-bound melanocortin 1 receptor (MC1R) (PDB code: 7F4D), somatostain-14-bound somatostatin receptor 2 (SSTR2) (PDB code: 7T10) and arginine-vasopressin (AVP)-bound vasopressin receptor 2 (V2R) (PDB code: 7DW9) **(e)**, the C terminus of peptides inserts into the TMD binding pocket such as angiotensin II-bound angiotensin II receptor type 1 (AT1R) (PDB code: 6OS0) and angiotensin II receptor type 2 (AT2R) (PDB code: 6JOD), Des-Arg10-kallidin bound (type 1 bradykinin receptor) B1R (PDB code: 7EIB), bradykinin-bound B2R (PDB code: 7F2O), cholecystokinin-8 (CCK-8)-bound $CCK_AR$ (PDB code: 7EZH) and gastrin-17 bound cholecystokinin B receptor ($CCK_BR$) (PDB code: 7F8V), JMV449-bound neurotensin receptor 1 (NTSR1) (PDB code: 6OS9), neuromedin U-bound neuromedin U receptor 2 (NUM2R) (PDB code: 7W55) and neuromedin S-bound NUM2R (PDB code: 7W57) **(f)**, and unique binding mode of galanin with galanin receptor 1 (GAL1R) (PDB code: 7WQ3) and galanin receptor 2 (GAL2R) (PDB code: 7WQ4) **(g).**

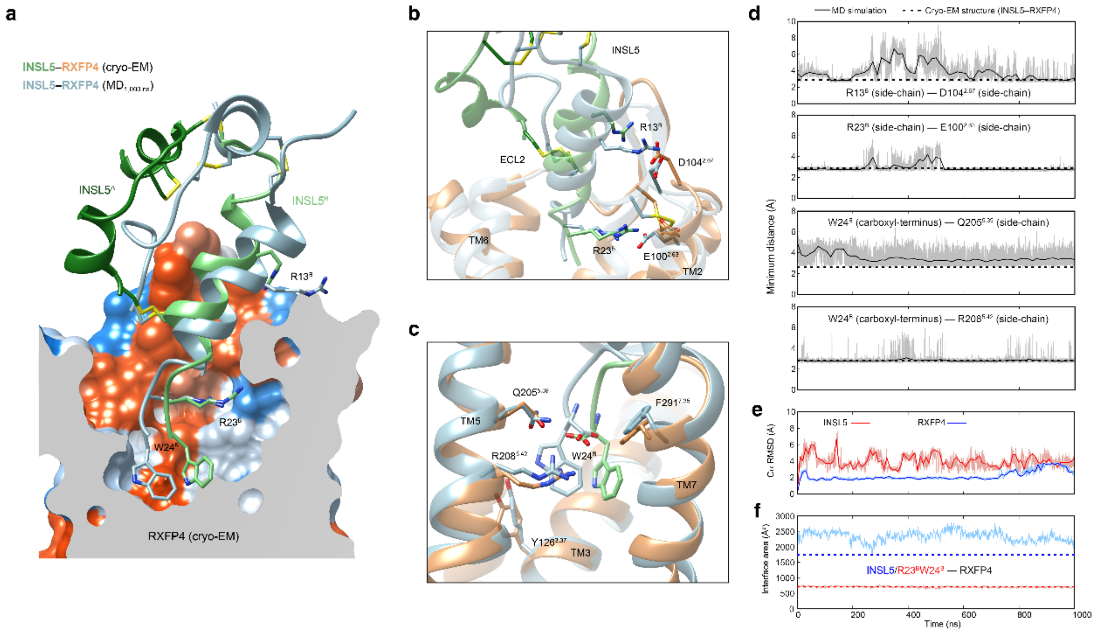

**Supplementary Figure 8. MD simulations of INSL5-bound active RXFP4. a,** Comparison of the INSL5 conformation between the final simulation snapshot at 1,000 ns and the cryo-EM structure of INSL5–RXFP4–$G_i$ complex. The key residues in the peptide-receptor interface are shown in sticks. **b-d,** Close-up views of the interactions between the C terminal α-helix of INSL5 B chain and receptor residues and their minimum distances during MD simulations. **e,** RMSD of Cα positions of the RXFP4 and INSL5, where all snapshots were superimposed on the cryo-EM structure of RXFP4 and INSL5 using the Cα atoms, respectively. **f,** The interface area between RXFP4 and INSL5 (blue) or the two C terminus residues $R23^B$ and $W24^B$ (red), calculated by freeSASA 2.0. The thick and thin traces represent moving averages and original, unsmoothed values obtained from one single MD simulation trajectory, respectively. The MD simulations were repeated independently three times with similar results.

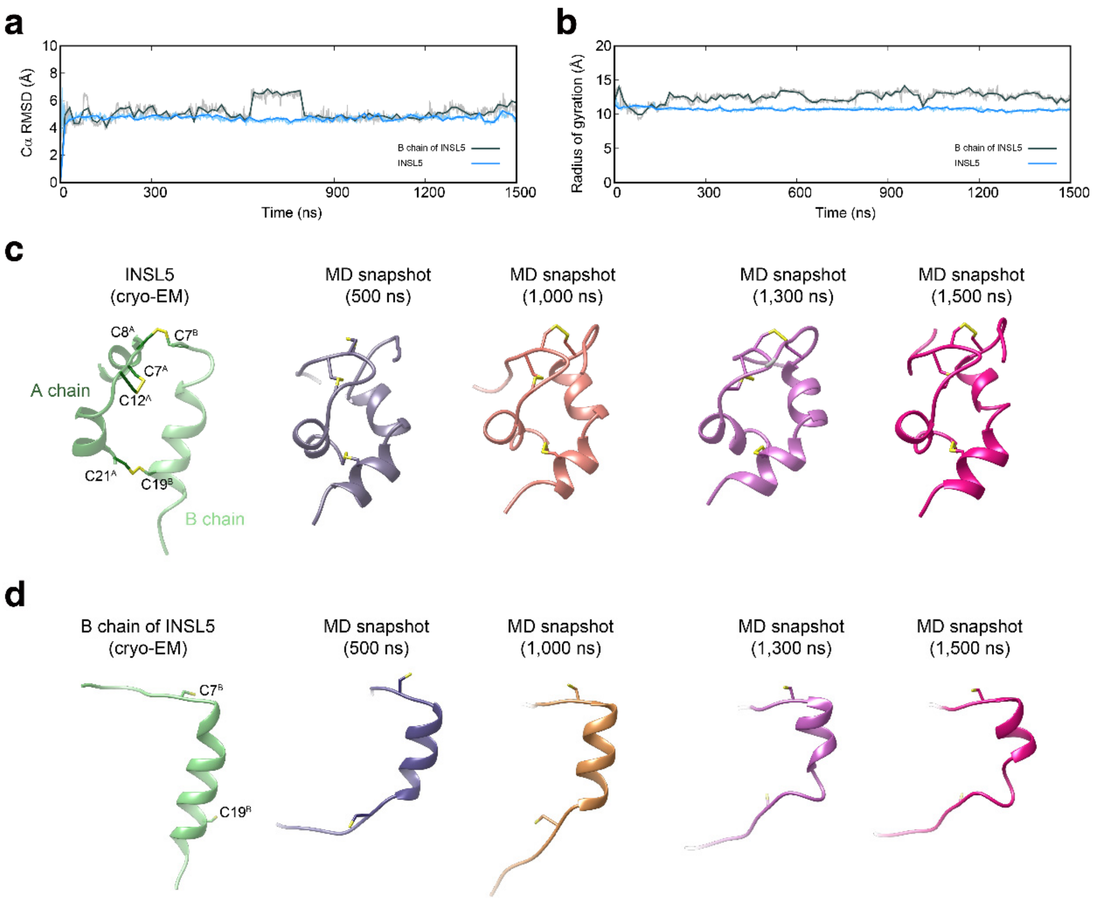

**Supplementary Figure 9. MD simulations of INSL5 and its B chain. a,** RMSD of Cα positions of the INSL5 and its B chain, where all snapshots were superimposed on the cryo-EM structure of INSL5 and its B chain using the Cα atoms, respectively. **b,** Radius of gyration of non-hydrogen atoms of the INSL5 and its B chain during MD simulations. **c,** Representative snapshots of MD simulations of INSL5. The cysteines are shown in sticks. **d,** Representative snapshots of MD simulations of the B chain of INSL5. The cysteines are shown in sticks. The thick and thin traces represent moving averages and original, unsmoothed values obtained from one single MD simulation trajectory, respectively. The MD simulations were repeated independently three times with similar results.

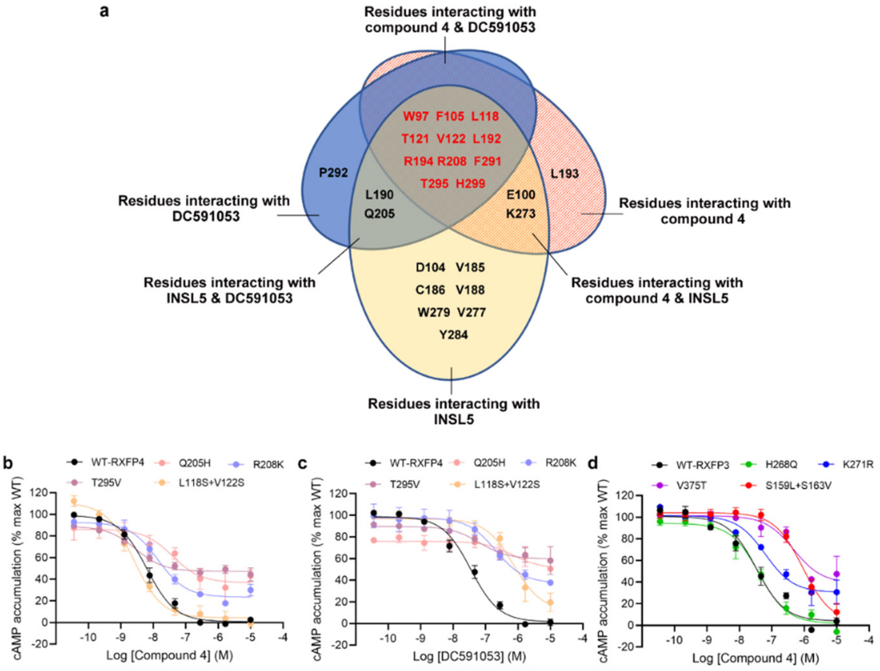

**Supplementary Figure 10. Peptidomimetic agonism and subtype selectivity of compound 4 and DC591053. a,** RXFP4 residues are categorized according to their interactions with the three ligands. **b-d,** Effects of amino acid switch in equivalent positions between RXFP4 and RXFP3 around the ligand-binding pocket on compound 4 **(b)** and DC591053 **(c)** induced cAMP accumulation in RXFP4-expressing cells as well as on compound 4 induced cAMP accumulation in RXFP3 **(d)**. Data are shown as means ± S.E.M. of at least three independent experiments.

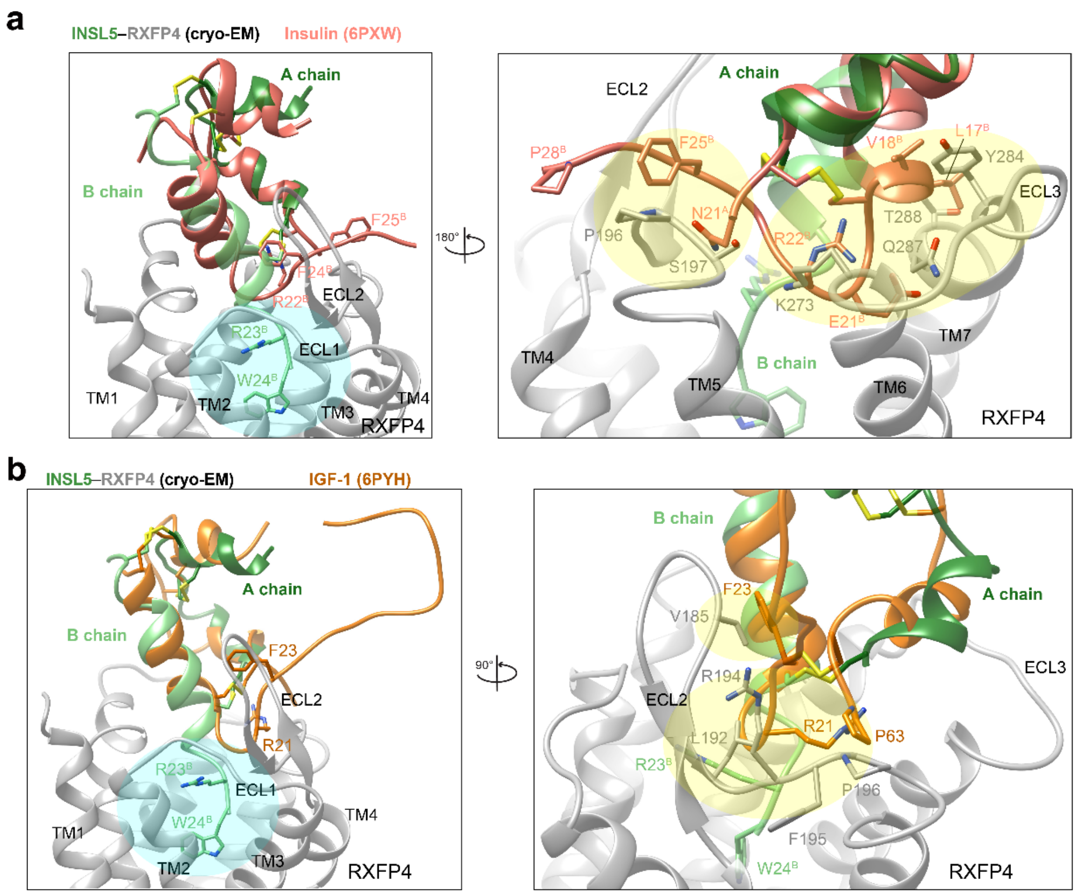

**Supplementary Figure 11. Superimposition of INSL5 from the INSL5–RXFP4–G$_i$ complex structure with insulin or IGF-1. a,** Insulin (salmon, PDB code: 6PXW) was superimposed on the INSL5 (green) from the cryo-EM structure of INSL5–RXFP4–G$_i$ using the Cα atoms. The three disulfide bonds and key residues in the peptide-receptor interface are shown in sticks. Compared to the endogenous agonist INSL5, the aligned insulin loses multiple potent interactions (cyan-shaded region) and causes significant steric clashes (yellow-shaded region) with RXFP4 (gray). **b,** IGF-1 (orange, PDB code: 6PYH) was superimposed on the INSL5 (green) from the cryo-EM structure of INSL5–RXFP4–G$_i$ using the Cα atoms. Compared to the endogenous agonist INSL5, the aligned IGF-1 loses multiple potent interactions (cyan-shaded region) and causes significant steric clashes (yellow-shaded region) with RXFP4 (gray).

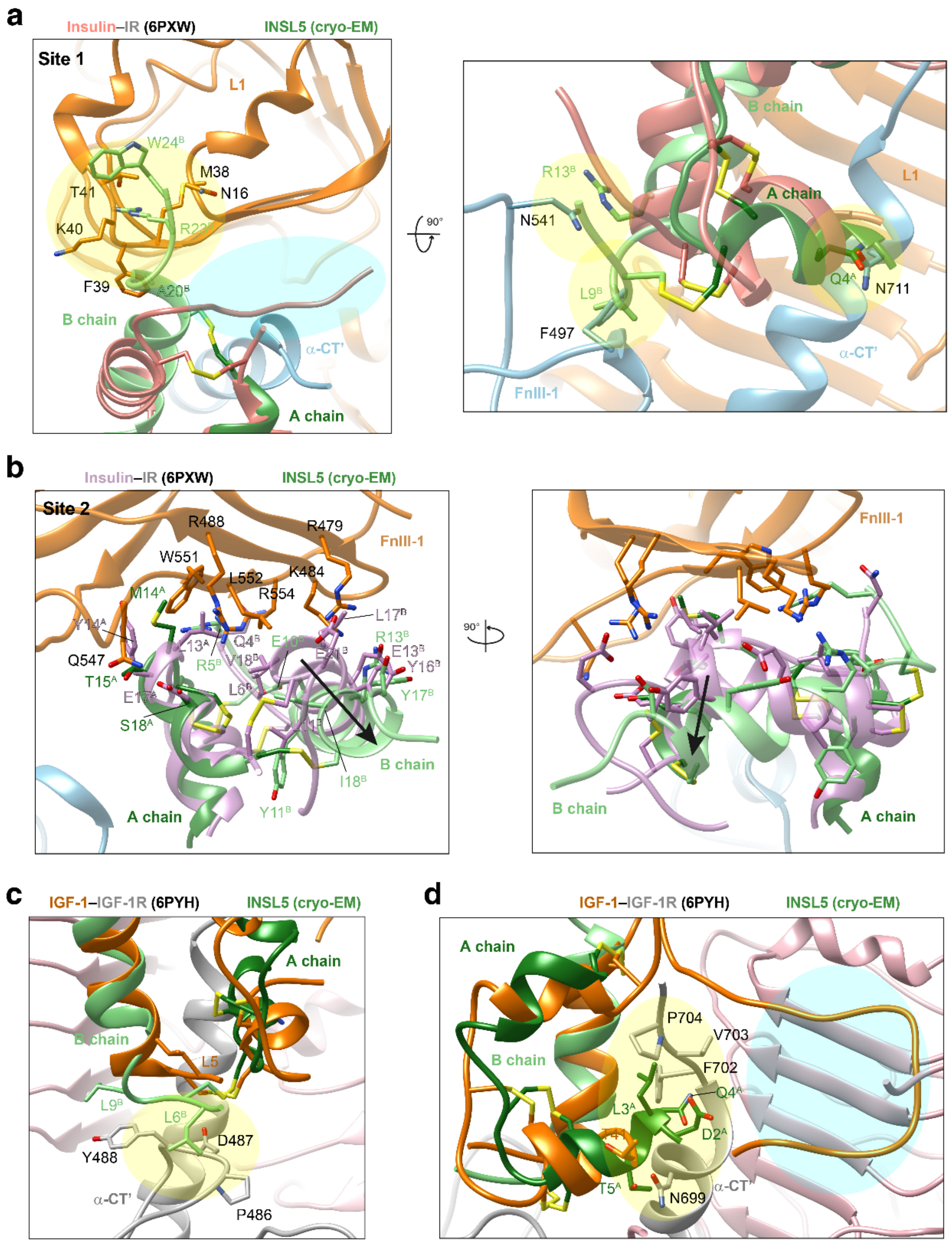

**Supplementary Figure 12. Superimposition of INSL5 to insulin or IGF-1 in complex with cognate receptors. a-b,** INSL5 (green) was superimposed on the insulin at site 1 (salmon, **a**) or at site 2 (plum, **b**) from the cryo-EM structure of insulin–insulin receptor (PDB code: 6PXW) using the Cα atoms. The three disulfide bonds and key residues in the peptide-receptor interface are shown in sticks. Compared to the endogenous agonist insulin, the aligned INSL5 loses multiple potent interactions (cyan-shaded region) and causes significant steric clashes (yellow-shaded region) with insulin receptor. **c,** INSL5 (green) was superimposed on the IGF-1 (orange) from the cryo-EM structure of IGF-1–IGF-1R (PDB code: 6PYH) using the Cα atoms. The three disulfide bonds and key residues in the peptide-receptor interface are shown in sticks. **d,** Compared to the endogenous agonist IGF-1, the aligned INSL5 loses multiple potent interactions (cyan-shaded region) and causes significant steric clashes (yellow-shaded region) with IGF-1R.

**Supplementary Table 1** Cryo-EM data collection, refinement and validation statistics.

| | **INSL5–RXFP4 (1-374)–$G_i$** | **Compound 4–RXFP4 (1-374)–$G_i$** | **DC591053–RXFP4 (1-374)–$G_i$** |
|---|---|---|---|
| **Data collection and processing** | | | |
| Magnification | 46,685 | 46,685 | 46,685 |
| Voltage (kV) | 300 | 300 | 300 |
| Electron exposure ($e^-/Å^2$) | 80 | 80 | 80 |
| Defocus range (μm) | -1.2 to -2.2 | -1.2 to -2.2 | -1.2 to -2.2 |
| Pixel size (Å) | 1.071 | 1.071 | 1.071 |
| Symmetry imposed | Cl | Cl | Cl |
| Initial particle images (no.) | 10,618,534 | 4,796,219 | 8,996,005 |
| Final particle images (no.) | 524,035 | 243,800 | 225,327 |
| Map resolution (Å) | 3.19 | 3.03 | 2.75 |
| FSC threshold | 0.143 | 0.143 | 0.143 |
| Map resolution range (Å) | 2.5-5.0 | 2.5-5.0 | 2.5-5.0 |
| **Refinement** | | | |
| Initial model used (PDB code) | PDB codes 7F2O and 6D9H | PDB codes 7F2O and 6D9H | PDB codes 7F2O and 6D9H |
| Model resolution (Å) | 3.3 | 3.2 | 3.1 |
| FSC threshold | 0.5 | 0.5 | 0.5 |
| Model resolution range (Å) | 2.5-5.0 | 2.5-5.0 | 2.5-5.0 |
| Map sharpening B factor ($Å^2$) | -163.30 | -96.90 | -106.50 |
| Model composition | | | |
| Non-hydrogen atoms | 8978 | 8687 | 8721 |
| Protein residues | 1154 | 1112 | 1116 |
| B factors ($Å^2$) | | | |
| Protein | 93.61 | 80.83 | 78.02 |
| Ligand | — | 115.88 | 175.71 |
| R.m.s. deviations | | | |
| Bond lengths (Å) | 0.004 | 0.004 | 0.004 |
| Bond angles (°) | 0.599 | 0.552 | 0.666 |

| | | | |
|---|---|---|---|
| Validation | | | |
| MolProbity score | 1.84 | 1.69 | 1.79 |
| Clash score | 9.24 | 8.75 | 8.03 |
| Poor rotamers (%) | 0.00 | 0.00 | 0.00 |
| Ramachandran plot | | | |
| Favored (%) | 94.96 | 96.61 | 94.98 |
| Allowed (%) | 5.04 | 3.39 | 5.02 |
| Disallowed (%) | 0.00 | 0.00 | 0.00 |

**Supplementary Table 2** Interactions of INSL5, compound 4 and DC591053 with RXFP4.

| RXFP4 | INSL5 | Compound 4 | DC591053 |
|---|---|---|---|
| W97$^{2.60}$ | Stacking | Stacking | Stacking |
| E100$^{2.63}$ | Salt bridge | Salt bridge | —— |
| D104$^{2.67}$ | Hydrogen bond | —— | —— |
| F105$^{ECL1}$ | Stacking | Stacking | Stacking |
| L118$^{3.29}$ | Hydrophobic contact | Hydrophobic contact | Hydrophobic contact |
| T121$^{3.32}$ | Hydrogen bond | Hydrophobic contact | Hydrophobic contact |
| V122$^{3.33}$ | Hydrophobic contact | Hydrophobic contact | Hydrophobic contact |
| V185$^{ECL2}$ | Hydrophobic contact | —— | —— |
| C186$^{ECL}$ | Hydrophobic contact | —— | —— |
| V188$^{ECL2}$ | Hydrophobic contact | —— | —— |
| L190$^{ECL2}$ | Hydrophobic contact | —— | Hydrophobic contact |
| L192$^{45.51}$ | Hydrophobic contact | Hydrophobic contact | Hydrophobic contact |
| L193$^{45.52}$ | —— | Hydrophobic contact<br>Hydrogen bond | —— |
| R194$^{ECL2}$ | Weak hydrogen bond | Hydrogen bond | Stacking |
| Q205$^{5.39}$ | Hydrogen bond | —— | Hydrogen bond |
| R208$^{5.42}$ | Salt bridge<br>Stacking | Stacking | Hydrogen bond<br>Stacking |
| K273$^{6.62}$ | Weak hydrogen bond<br>Hydrophobic contact | Hydrogen bond | —— |
| V277$^{ECL3}$ | Hydrophobic contact | —— | —— |
| W279$^{ECL3}$ | Hydrophobic contact | —— | —— |
| Y284$^{7.28}$ | Hydrophobic contact | —— | —— |
| F291$^{7.35}$ | Stacking | Stacking | Stacking |
| P292$^{7.36}$ | —— | —— | Hydrophobic contact |
| T295$^{7.39}$ | Hydrophobic contact | Hydrogen bond | Hydrogen bond |
| H299$^{7.43}$ | Stacking | Hydrogen bond | Stacking |

**Supplementary Table 3.** Ligand-mediated inhibition of forskolin-induced cAMP accumulation.

| **Receptor** | **INSL5** | | **Compound 4** | | **DC591053** | |
|---|---|---|---|---|---|---|
| | $pEC_{50}$ ± S.E.M. | $E_{max}$ ± S.E.M. (% WT) | $pEC_{50}$ ± S.E.M. | $E_{max}$ ± S.E.M. (% WT) | $pEC_{50}$ ± S.E.M. | $E_{max}$ ± S.E.M. (% WT) |
| RXFP4-WT | 9.07 ± 0.07 | 99.76 ± 2.15 | 7.94 ± 0.08 | 99.99 ± 2.85 | 7.68 ± 0.09 | 99.14 ± 2.97 |
| HA-H10-Bril-RXFP4-15AA-LgBiT | 8.94 ± 0.08 | 110.76 ± 2.64 | 7.74 ± 0.07 | 110.40 ± 2.36 | 7.14 ± 0.08 | 108.57 ± 2.51 |

All data were fitted with a three-parameter logistic curve to obtain $pEC_{50}$ and $E_{max}$ values. The assay was performed in transiently transfected HEK293T cells. Data represent means ± S.E.M. of at least three independent experiments performed in quadruplicate. Statistical analysis was performed using a two-tailed Student's *t*-test and no significance was found among the values. WT, wild-type.

**Supplementary Table 4.** Effects of residue mutation on ligand-mediated inhibition of forskolin-induced cAMP accumulation.

| Receptor | Mutation | INSL5 | | Compound 4 | | DC591053 | |
|---|---|---|---|---|---|---|---|
| | | $p$EC$_{50}$ ± S.E.M. | E$_{max}$ ± S.E.M. (% WT) | $p$EC$_{50}$ ± S.E.M. | E$_{max}$ ± S.E.M. (% WT) | $p$EC$_{50}$ ± S.E.M. | E$_{max}$ ± S.E.M. (% WT) |
| RXFP4 | WT | 9.12 ± 0.06 | 99.88 ± 2.28 | 8.20 ± 0.07 | 98.22 ± 3.01 | 7.46 ± 0.07 | 97.18 ± 3.11 |
| | W97A | 7.81 ± 0.18*** | 61.33 ± 5.34**** | N.A. | N.A. | 7.25 ± 0.27 | 108.65 ± 10.04 |
| | E100A | N.A. | N.A. | N.A. | N.A. | 6.97 ± 0.18 | 79.27 ± 6.72 |
| | D104A | 8.79 ± 0.25 | 59.09 ± 6.41**** | N.D. | N.D. | N.D. | N.D. |
| | F105A | 8.00 ± 0.11** | 80.16 ± 4.08 | 7.81 ± 0.21 | 60.11 ± 5.70**** | 6.77 ± 0.15 | 57.39 ± 4.17**** |
| | T121A | N.A. | N.A. | 6.90 ± 0.30**** | 59.77 ± 8.29**** | 6.14 ± 0.41** | 31.53 ± 7.09**** |
| | R194A | 8.78 ± 0.17 | 79.76 ± 6.06 | 7.33 ± 0.24 ** | 78.96 ± 8.75 | 6.55 ± 0.19* | 81.10 ± 7.60 |
| | Q205A | 8.40 ± 0.30 | 50.94 ± 6.54**** | N.D. | N.D. | N.D. | N.D. |
| | R208A | N.A. | N.A. | 7.38 ± 0.17** | 63.76 ± 4.94*** | 6.58 ± 0.16* | 52.09 ± 4.26**** |
| | K273A | 8.45 ± 0.29 | 58.10 ± 7.38**** | 7.33 ± 0.19** | 76.06 ± 6.54** | 5.73 ± 0.18**** | 68.51 ± 8.34** |
| | W279A | 8.73 ± 0.17 | 43.30 ± 3.29**** | N.D. | N.D. | N.D. | N.D. |
| | Y284A | 8.74 ± 0.19 | 49.25 ± 4.06**** | N.D. | N.D. | N.D. | N.D. |
| | H299A | 9.02 ± 0.29 | 30.77 ± 3.77**** | N.A. | N.A. | 8.44 ± 0.24* | 36.04 ± 4.01**** |
| | L118S + V122S | N.D. | N.D. | 8.60 ± 0.10 | 106.23 ± 4.75 | 6.14 ± 0.18*** | 84.85 ± 8.36 |
| | Q205H | N.D. | N.D. | 7.41 ± 0.32* | 49.35 ± 7.02**** | 6.17 ± 0.46*** | 25.10 ± 6.36**** |
| | R208K | N.D. | N.D. | 7.85 ± 0.13 | 69.09 ± 4.02** | 6.69 ± 0.22 | 60.43 ± 6.52*** |
| | T295V | N.D | N.D. | 8.55 ± 0.15 | 41.58 ± 2.93**** | 7.19 ± 0.33 | 30.00 ± 4.54**** |
| RXFP3 | WT | N.D. | N.D. | 7.48 ± 0.12 | 96.70 ± 5.12 | N.D. | N.D. |
| | S159L + S163V | N.D. | N.D. | 6.05 ± 0.12*** | 101.29 ± 6.72 | N.D. | N.D. |
| | H268Q | N.D. | N.D. | 7.39 ± 0.11 | 92.52 ± 4.71 | N.D. | N.D. |
| | K271R | N.D | N.D. | 7.21 ± 0.17 | 70.86 ± 5.48**** | N.D | N.D. |
| | V375T | N.D. | N.D. | 6.33 ± 0.24** | 62.11 ±7.78**** | N.D | N.D. |

Inhibition of forskolin-induced cAMP accumulation was performed in HEK293T cells transiently transfected with WT and mutant receptors. All the mutant constructs were modified by single-point mutation in the setting of the WT receptor. cAMP accumulation data were analyzed using a three-parameter logistic equation to determine $p$EC$_{50}$ and E$_{max}$ values. E$_{max}$ values for mutants are defined as the window between the maximal response and vehicle control (no ligand) and expressed as a percentage of the WT. Data shown are means ± S.E.M. of at least three independent experiments performed in quadruplicate. One-way ANOVA were used to determine statistical difference (*P< 0.05, **P< 0.01, ***P< 0.001 and ****P< 0.0001). N.A., not active; N.D., not determined.
